# 4-methyl-3-aminopyridine: A novel active blocker of voltage-gated potassium ion channels in the central nervous system

**DOI:** 10.64898/2026.03.06.710137

**Authors:** Sofia Rodríguez-Rangel, Oscar Gutiérrez Coronado, Brenda Mata Ortega, Yang Sun, Saadi El-Saadi, Pedro Brugarolas, Jorge E. Sánchez-Rodríguez

**Affiliations:** Departamento de Física, Universidad de Guadalajara, CUCEI, Blvd. Marcelino García Barragán 1421, C.P. 44430, Guadalajara, Jalisco, México; Departamento de Ciencias de la Tierra y de la Vida, Universidad de Guadalajara. CULagos, Av. Enrique Diaz de León 1144, C.P. 47463, Lagos de Moreno, Jalisco, México; Department of Radiology, Massachusetts General Hospital and Harvard Medical School, Boston, MA, 02114, USA

**Keywords:** 4-Aminopyridine, 4-methyl-3-aminopyridne, Potassium ion channel blocker, Electrophysiology, Acute toxicity and Pharmacokinetics

## Abstract

Aminopyridines, including 4-aminopyridine (4AP), 3,4-diaminopyridine, and [^18^F]3-fluoro-4-aminopyridine, are voltage-gated potassium (K_V_) channel blockers used clinically to enhance conduction in neurological disorders and to image demyelination by PET. Developing new aminopyridines may yield improved therapeutics or imaging agents. Here, we characterized the physicochemical properties (*pK*_*a*_, log *D*), K_V_ channel–blocking activity, toxicity (*LD*_50_), and pharmacokinetics of a novel compound, 4-methyl-3-aminopyridine (4Me3AP). 4Me3AP was less basic and more lipophilic than 4AP and showed greater blocking potency across multiple K_V_ channels expressed in *Xenopus* oocytes. In mice, 4Me3AP exhibited lower acute toxicity (*LD*_50_= 29.3 mg/kg) than 4AP (*LD*_50_= 12.7 mg/kg) and a longer plasma half-life. These findings indicate that 4Me3AP is a potent K_V_ channel blocker with favorable pharmacological properties, supporting its potential for symptomatic treatment of demyelinating diseases.

## Introduction

Aminopyridines are a class of small heterocyclic compounds that act as voltage-gated potassium (K_V_) channel blockers and have long been used to enhance conduction in demyelinated axons^1–3^. The prototypical compound, 4-aminopyridine (4AP), is FDA-approved for improving walking in people with multiple sclerosis (MS)^4^ and has shown benefit across a range of central nervous system (CNS) demyelinating conditions by prolonging action potentials, restoring conduction in partially demyelinated fibers, and enhancing synaptic transmission^1,5,6^. In contrast, 3,4-diaminopyridine (3,4DAP) is primarily used to treat peripheral nervous system (PNS) disorders, such as Lambert–Eaton myasthenic syndrome, where it increases acetylcholine release at the neuromuscular junction and improves muscle strength^7–9^. Beyond their therapeutic applications, radiolabeled aminopyridines, such as [^18^F]3-fluoro-4-aminopyridine, enable PET imaging of demyelination by binding K_V_ channels that become exposed and upregulated following myelin loss^10,11^. Because the potency, safety, and pharmacokinetic behavior of aminopyridines depend strongly on their physicochemical properties and their interactions with specific Kv channel subtypes^12–15^, developing new derivatives offers promising opportunities to improve both therapy and imaging.

The mechanisms and structure–activity relationships (SAR) of aminopyridines are strongly governed by their acid–base and lipophilic properties^7,12–16^. Aminopyridines exist in a *pH*-dependent equilibrium between a protonated (charged) form and a neutral (uncharged) form, and the ratio of these species is determined by the compound’s *pK*_*a*_. This equilibrium is functionally crucial: the *protonated* form is required for binding and blocking Kv channels from the intracellular side^17–20^, whereas the *neutral* form is necessary for passive diffusion across lipid membranes, including the blood–brain barrier (BBB)^7,11,13,14,21^. Thus, *pK*_*a*_ directly influences both channel-blocking potency and CNS accessibility by dictating the fraction of molecules available in each state at physiological *pH*. Lipophilicity (log *D*) further modulates membrane permeability, tissue distribution, and whole-body pharmacokinetics, including plasma half-life and clearance^22,23^. Small structural modifications to the aminopyridine scaffold, such as methylation or fluorination, can shift *pK*_*a*_ altering the protonated/neutral ratio, change lipophilicity, and thereby modify K_V_ channel binding affinity, BBB permeability, and systemic pharmacokinetics. These SAR principles guide the rational design of next-generation aminopyridines with optimized therapeutic or imaging profiles.

Despite the therapeutic and imaging value of this class, nearly all known aminopyridine derivatives, including clinical drugs and PET tracers, are based on the 4AP scaffold, with the amino group positioned at the 4-ring position. However, the clinical efficacy of 3,4DAP raises the possibility that compounds derived from 3-aminopyridine (3AP) may also possess significant biological activity as Kv channel blockers, an area that has not been explored for single-amino aminopyridines. In this study, we describe and characterize several novel aminopyridines in the context K_V_ channel–blocking activity including 4-methyl-3-aminopyridine (4Me3AP), the first 3-aminopyridine derivative to show high blocking potency. Furthermore, we report the physicochemical, pharmacological, toxicological, and pharmacokinetic properties of this compound to demonstrate its potential as a novel aminopyridine with therapeutic relevance.

## Materials and Methods

### Materials

All study compounds were purchased from Sigma-Aldrich or Chem-Impex International. A full table of key resources and materials can be found in the Supplementary Material **Table S1**.

### *pK*_*a*_ determination

The *pK*_*a*_ for each 4AP-based structural analog was estimated from titration curve data by titrating an initial solution of 10 mM of each compound with HCl at 0.01 M in a water type I (Milli Q). The recorded *pH* was plotted as a function of the volume of HCl added. Subsequently, the *pK*_*a*_was determined at the halfway point to the equivalence point of the titration curve. This procedure was repeated at least 4 times for each compound.

### Octanol-water partition coefficient determination

The octanol-water partition coefficient (log *D*) of each 4AP-based structural analog was assessed using the shake flask method. To this end, 3 μL at 200 mM of each compound was added to 3 mL of Phosphate Buffered Saline solution (PBS, Gibco, Thermo Fisher Scientific Inc.) at *pH* 7.4 and 3 mL of 1-octanol (Sigma-Aldrich, Merck KGaA, Darmstadt, Germany) and partitioned via vortexing. The mixture was placed in a separatory funnel until the phases were separated and homogenized. Subsequently, the relative concentration of each 4AP-based structural analog partitioned in each phase was measured by determining its absorbance by standard procedures of UV-VIS spectroscopy. A calibration curve of absorbance *vs.* concentration for each compound within the linear range (typically 0–50 μM) was constructed prior to the phase separation procedure. This procedure was repeated at least 4 times for each compound.

### Permeability rate determination

The permeability rates (*P*_*e*_) of the compounds were determined using parallel artificial membrane permeability assay-blood–brain barrier (BBB) kit (BioAssay systems, Hayward, USA) following the manufacturer’s protocol. Initially, solutions of each test compound were prepared in DMSO at a concentration of 10 mM. These stock solutions along with the stock solutions of control compounds (high control: promazine hydrochloride, low control: diclofenac) were then diluted with PBS (*pH* = 7.2) to obtain the donor solutions with a final concentration of 500 μM. At the same time, 200 μM of equilibrium standards for each compound and a DMSO blank control solution were prepared. In the experimental setup, 300 μL of PBS was added to the desired well of the acceptor plate, and 5 μL of BBB lipid solution in dodecane was added to membranes of the donor plate. Next, 200 μL of the donor solutions of each test compound and each permeability control were added to the duplicate wells of the donor plate. The donor plate was carefully placed on the acceptor plate and incubated for 18 h at room temperature. After incubation, UV absorption measurements were conducted using 100 μL of the resulting solutions from the acceptor plate and the equilibrium standards. UV absorption of the controls was measured by running a UV scan in the range of 200–500 nm. UV absorption of the compounds was measured using HPLC equipped with a UV detector and C18 column.

### Synthesis of K_V_ ion channels RNA

RNAs used in the present study were synthesized *in vitro* by standard procedures of Molecular Biology. The cDNA clones encoding for the Kv ion channel (gen, vector, restriction enzyme, promoter, and specie) used in this study were: *i*) Shaker K_V_ (Shaker, pGSTA, Not I, T7, *Drosophila melanogaster*), *ii*) Kv 1.2 (KCNA2, pMAX, Pme I, T7, *Rattus norvegicus*), *iii*) Kv1.3 (KCNA3, pGSTA, Not I, T7, *Homo sapiens*), *iv*) K_V_ 2.1 (KCNB1, pGEM-HE, Nhe I, T7, *Homo sapiens*), *v*) K_V_ 6.4 (KCNG4, pGEM-HE, Nhe I, T7, *Homo sapiens*), *vi*) K_V_ 7.2 (KCNQ2, pTLN, MIu I, SP6, *Homo sapiens*), *vii*) K_V_ 7.3 (KCNQ3, pTLN, Hpa I, SP6, *Homo sapiens*), *viii*) K_V_ 10.1 (KCNH1, pSGEM, Nhe I, T7, *Homo sapiens*) and *ix*) K_V_ 11.1 (KCNH2, pSP64, EcorI, SP6, *Homo sapiens*). Each cDNA was amplified via a miniprep preparation kit (QIAGEN GmbH, Germany) and then linearized with the indicated restriction enzyme (New England Biolabs, Inc. Ipswich, MA, USA). Subsequently, each RNA was synthesized using the mMESSAGE cRNA transcription kit (AmbionTM, MEGAscript®, Austin, TX, USA) according to the specified promoter.

### Expression of K_V_ ion channels in Xenopus laevis oocytes

Each K_V_ ion channel was heterologously expressed in *Xenopus laevis* oocytes. To this end, only mature *Xenopus laevis* frogs (Xenopus 1, Corp., MI., USA, and Aquanimals SA de CV, Queretaro, Mexico) held in captivity were used as oocyte suppliers by extracting a volume of ∼1 mL from the ovarian lobes via survival surgery under anesthesia with MS-222. Subsequently, oocytes were isolated under the simultaneous process of the enzymatic treatment of collagenase type II (Worthington Biochemical Corp., NJ, USA) and mechanical agitation. Then, each isolated oocyte was injected with 15-25 ng of RNA encoding for each K_V_ by using the Nanoinyect II Auto-Nanoliter Injector system (Drummond Scientific Company, Broomall, USA) and incubated before the experiments from 1 to 5 days in a Standard Oocytes Saline (SOS) solution containing (in mM): 100 NaCl, 1 MgCl2, 10 HEPES, 2 KCl and 1.8 CaCl2 with 50 μg/mL gentamycin at *pH* 7.5 in an atmosphere at 17 °C. For the expression of the heteromeric K_V_ 7.2/K_V_ 7.3 or K_V_ 2.1/K_V_ 6.4 ion channels, each oocyte was co-injected with the corresponding cRNA with a stoichiometry of 1:1 or 3:1, respectively^24,25^.

### Cut-Open Voltage Clamp (COVC) Electrophysiology and blocking potency of 4AP-based structural analogs

Blocking potency of 4AP-based structural analogs was evaluated in terms of the half-maximal inhibitory concentration (*IC*_50_) needed to inhibit 50% of K^+^ current yielded by the Shaker Kv ion channel and other K_V_ ion channels expressed in *Xenopus laevis* oocytes and under voltage-clamp conditions with COVC methodology as previously described^11–13^. For COVC procedures, internal and external solutions were composed (in mM) of: 120 KOH, 2 EGTA, 20 HEPES and 12 KOH, 2 Ca(OH)_2_, 105 NMDG (N-methyl-D-glucamine)-methylsufonate (MES), 20 mM HEPES, respectively. Both solutions were adjusted at *pH* 7.4. For measurements where the *pH* was varied, HEPES was interchanged by 2-(cyclohexylamino)ethanesulfonic acid (CHES, pH=9.1) or 2-(N-morpholino)ethanesulfonic acid hydrate (MES-hydrate, pH=5.7 or 6.8) buffers. For *IC*_50_determinations, a relative current (*I*_*r*_) *vs*. the concentration of each 4AP-based structural analog ([*X*]) a curve was measured. To this end, K^+^ current was recorded first in the absence of the inhibitor (*I*) from each oocyte expressing a K_V_ ion channel as response of membrane depolarization with a voltage stimulus protocol that consisted in steps of 50 ms (for Shaker K_V_ ion channel or > 50 ms (as indicated for other K_V_ channels) from −100 to 60 mV in increments of 10 mV. Subsequently, the current was measured in the presence of the inhibitor (*I*_*X*_) at sequential [*X*] values from 0.0001 to 10 mM. Ionic currents were amplified and digitized with the Oocyte Clamp Amplifier CA-1A (Dagan Corporation, Minneapolis, MN, USA) and the USB-1604-HS-2AO Multifunction Card (Measurement Computing, Norton, MA, USA), respectively, and controlled with the GpatchMC64 program (Department of Anesthesiology, UCLA, Los Angeles, CA, USA) via a PC. Electrical signals were sampled at 100 kHz and filtered at 10 kHz. All the experiments were performed at room temperature (21–23 °C).

### Electrophysiology data analysis

Electrophysiological recordings were analyzed using Clampfit (Axon Instruments), Analysis (Dept. of Physiology, UCLA), and OriginPro 8 (®OriginLab Corporation) software as previously described^12,13^. *IC*_50_ for each 4AP-based compound was determined by fitting the *I*_*r*_ (= *I*_*X*_⁄*I*) *vs.* [*X*] curve to the pharmacological Hill equation:

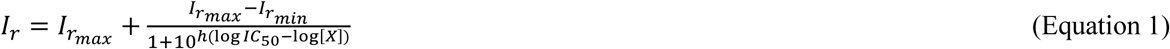

where *I*_*rmax*_ and *I*_*rmin*_ are the maximal and minimal value of *I*_*r*_ and ℎ is the Hill coefficient which value was typically in the range 0.9 < ℎ < 1.1. Voltage dependence of *IC*_50_(*IC*_50_(*V*)) was analyzed by fitting the *IC*_50_(*V*) curve with the one-step model of inhibition which allow to determine the fractional electrical-distance (δ) that each 4AP analog cross through the electric field generated in the K_V_ pore channel to reach its binding site^12,13,26^:

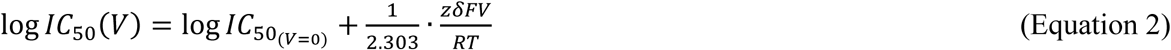

where *IC*_50(*V*=0)_ is the model-value of *IC*_50_ at *V* = 0 mV, *F* and *R* are the Faraday and universal gas constants, respectively, *T* is the temperature, and *Z* is the apparent charge.

### Acute toxicology assays

Acute toxicity study was carried out as follows. Each dose of 4AP, 3Me4AP and 4Me3AP was evaluated in independent groups of five male specimens of the *BALB/c* strain between 6 and 8 weeks of age with a weight of 20-25 g obtained from the Animal Facility of the Centro de Investigación Sobre Enfermedades Infecciosas, Instituto Nacional de Salud Pública (Cuernavaca, Morelos, Mexico). A dose in the range of 2.5 to 20 mg/kg (or 50 mg/kg of 4Me3AP) was administered to each mouse. The effects of each molecule were monitored for a period of 180 min. No macroscopic changes were observed in the organs of the mice treated with the different doses of 4AP, 3Me4AP and 4Me3AP. The acute oral median lethal dose (*LD*_50_) was determined by the Bliss method or Probits model^27^ after the administration of each dose.

### Pharmacokinetics procedures

The pharmacokinetics of 4AP and 4Me3AP molecules were evaluated in the murine *BALB/c* strain model. Prior to the study, the mice underwent a quarantine period and were subsequently housed at the Animal Facility of the Research and Innovation Building at the Universidad de Guadalajara’s Centro Universitario de los Lagos. To quantify a reliable concentration of 4AP and 4Me3AP in plasma, a single dose of 10 mg/kg of 4AP or 4Me3AP diluted in 200 µL of type I water was administered in each mouse using the gavage method. Subsequently, the plasma concentration of 4AP or 4Me3AP extracted from each mouse was determined at different time points within a period of 0 to 180 minutes. For this purpose, 10 minutes prior to blood sample extraction, each mouse was subcutaneously administered with a dose of 30 μL of Sodium Heparin (1000 IU) as an anticoagulant. Later, approximately 1.5 mL of blood sample was extracted via cardiac puncture under anesthesia. The solid and liquid components of the sample were separated by centrifugation for 10 minutes at 10,000 rpm and ambient temperature. Subsequently, the proteins present in the serum were precipitated by adding three volumes of High-Performance Liquid Chromatography (HPLC) –grade 100% acetonitrile for each volume of serum obtained according to the method proposed by Polson^28^. The mixture was centrifuged for 10 minutes at 10,000 rpm at 4°C. Finally, the supernatant was filtered and transferred to vials for HPLC analysis. The HPLC equipment used for this study was the ACQUITY Arc (Waters Corporation, Milford, Massachusetts, USA), equipped with the 2998 Photodiode Array (PDA) detector. The HPLC method implemented consisted of: mobile phase (12mM Na_2_HPO_4_, 8mM NaOH, 4% (m/v) MeCN. pH=12), stationary phase (XBridge BEH C18 Column, 130Å, 3.5 μm, 4.6 mm X 100 mm. (Waters Corporation, Milford, Massachusetts, USA)) with a flow rate (in mL/min) of 0.4, sample temperature (in °C) of 22, and column temperature (in °C) of 45, respectively. This method yielded a retention time of 4.9 min for 4AP and 13.6 min for 4Me3AP, quantified at 240 nm and 231 nm, respectively (**Figure S1**).

### Pharmacokinetic data analysis

Plasma concentrations of 4AP or 4Me3AP were determined over a 0–180 min time interval following administration by quantification of the corresponding chromatographic peak areas using an external calibration curve (**Figure S1**), in accordance with standard procedures implemented in the ACQUITY Arc driver software. Then, the pharmacokinetic parameters determined from the fitted concentration *vs.* time curve included the maximum plasma concentration (*C*_*max*_), time to reach maximum concentration (*t*_*max*_), elimination half-life (*t*_1⁄2_), elimination rate constant (*K*_*e*_), area under the concentration *vs.* time curve (*AUC*_0−250_), apparent volume of distribution (*V*_*d*_), and clearance (*CL*). For this purpose, the elimination phase of the pharmacokinetic curve for each drug was fitted using a simple pharmacokinetic model^29^ consisting of *m* compartments of the form:

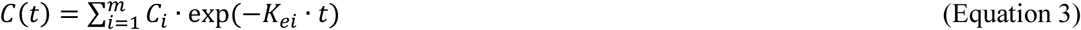

where *C*_*i*_ and *K*_*ei*_ are the drug concentration at *t* = 0 min and the drug elimination rate constant in the *i*-th compartment, respectively.

### Animal Studies Compliance

Methods involving *Xenopus laevis* frogs and *BALB/c* mice were performed in accordance with relevant guidelines and regulations and with the approval of the Comité Institucional del Cuidado y Uso de Animales en el Laboratorio at the University of Guadalajara, protocols: CUCEI/CINV/CICUAL-03/2023 and CUCEI/CINV/CICUAL-02/2023, respectively.

## Results

### Physicochemical and pharmacological characterization of 4-aminopyridine structural analogs

We sought to investigate commercially available pyridine derivatives with −NH_2_ or −CH_3_ substitution in 2-, 3– and 4– positions. The study compounds included the ten structurally related compounds shown in **Figure 1**: 2,4-diaminopyridine (2,4DAP), 3,4-diaminopyridine (3,4DAP), 2-methyl-4-aminopyridine (2Me4AP), 3-methyl-4-aminopyridine (3Me4AP), 4-methylpyridine (4MeP), 2,4-dimethylpyridine (2,4DMeP), 3,4-dimethylpyridine (3.4DMeP), 2-amino-4-methylpyridine (4Me2AP), 3-amino-4-methylpyridine (4Me3AP). To our knowledge, this is the first time 4MeP, 2,4DMeP, 3,4DMeP, 4Me2AP and 4Me3AP compounds are investigated in the context of K^+^ channel blocking activity.

**Figure 1.**
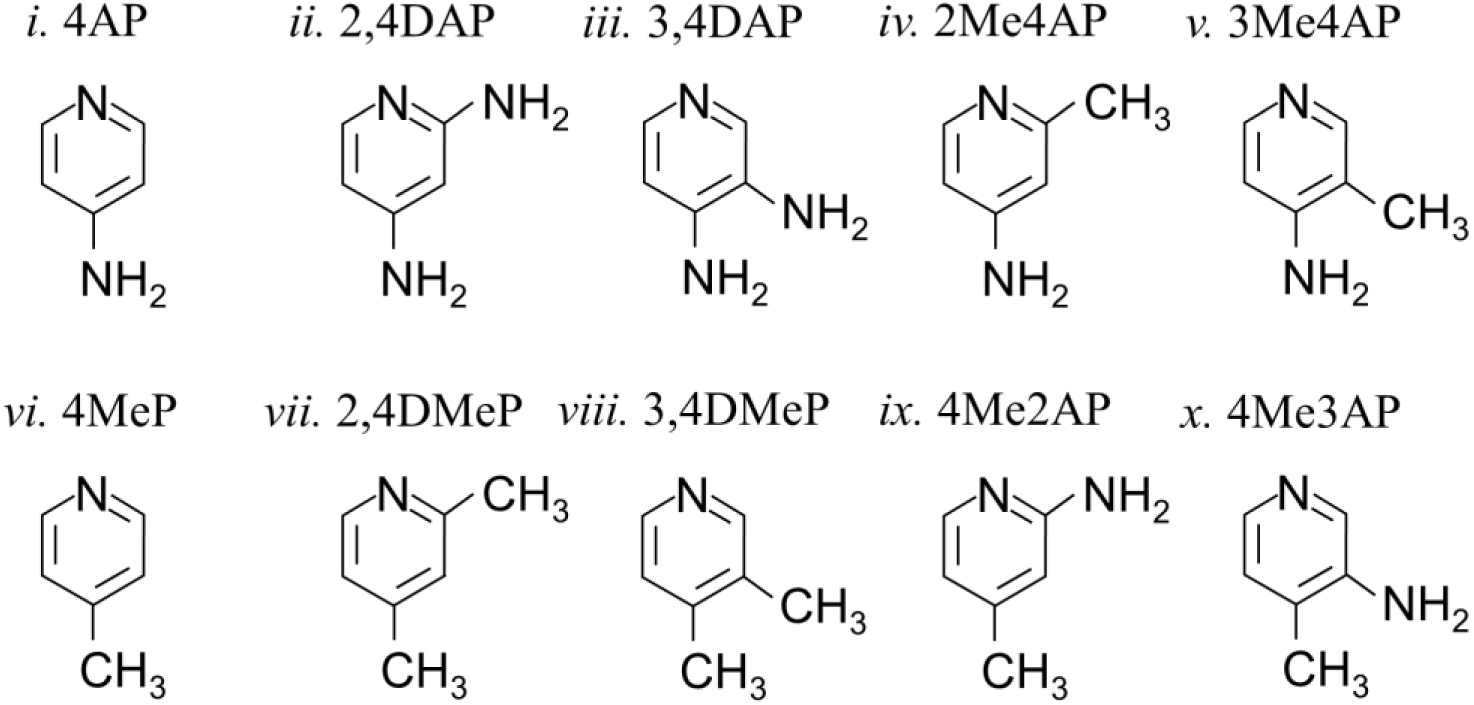
Chemical structures of 4-aminopyridine-based structural analogs. Structural analogs of 4-aminopyridine: i. 4-aminopyridine (4AP), ii. 2,4-diaminopyridine (2,4DAP), iii. 3,4-diaminopyridine (3,4DAP), iv. 2-methyl-4-aminopyridine (2Me4AP), v. 3-methyl-4-aminopyridine (3Me4AP), vi. 4-methylpyridine (4MeP), vii. 2,4-dimethylpyridine (2,4DMeP), viii. 3,4-dimethylpyridine (3,4DMeP), ix. 4-methyl-2-aminopyridine (4Me2AP), x. 4-methyl-3-aminopyridine (4Me3AP).

First, we evaluated the blocking potency of these compounds on the Shaker K_V_ ion channel, a functional model of the KCNA protein family, expressed in *Xenopus laevis* oocytes (**Figure 2**). **Figure 2a** shows representative K^+^ currents triggered as the response to a +40 mV voltage membrane stimulus on each oocyte expressing Shaker K_V_, before and after the addition of 1mM of each compound. Then, we quantitatively assessed the inhibition of the K^+^ current due to the sequential increase in compound concentration from 0.0001 to 10 mM for each compound in order to determine dose-response curves (**Figure 2b**). Subsequently, *IC*_50_values were determined by fitting each independent measured curve to the Hill pharmacological model (**Equation 1**). *IC*_50_ values are reported on the **Table I**. Compared to the *IC*_50_ value determined for 4AP (292 μ*M*),

**Figure 2.**
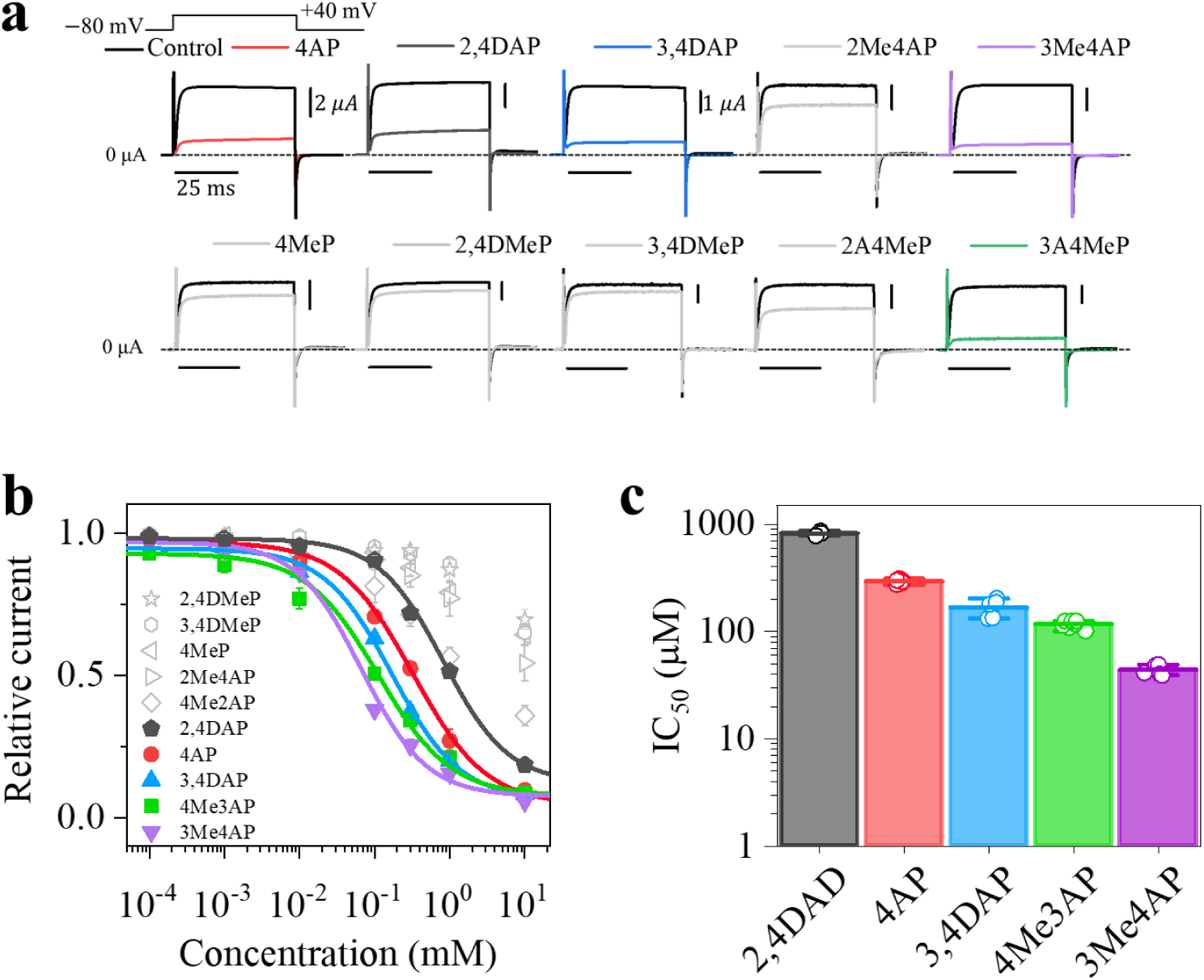
Chemical and pharmacological characterization of 4AP-based structural analogs. (**a**), Representative K^+^ recordings elicited from ten different oocytes expressing the Shaker K_V_ ion channel in response to a voltage stimulus (upper voltage protocol) before (upper, black) and after (lower, colored) addition of 1 mM of the external oocyte membrane vestibule of each 4AP-based structural analog. Dashed lines represent the zero-current level and horizontal and vertical bars indicate the time and current scale for each recording. (**b**), Relative current *vs.* the concentration curves of each 4AP-based structural analog evaluated at +40 mV and *pH* = 7.4. Continuous lines represent the fits with the Hill equation (**Equation 1**) with a Hill coefficient ranging from 0.9 to 1.1. (**c**), *IC*_50_values of the 4AP-based structural analogs determined from the Hill equation fitting of the data showed in panel b. The number of independent experiments (*n*) ranged from 3 to 7, and results are expressed as the mean ± standard deviation.

**Table I.** *pK*_*a*_ log *D* and *P*_*e*_ values of 4AP structural analogs.

| Drug | $pK_a$ | n | $\log D^*$ | n | $P_e$ (cm/s) | n |
| --- | --- | --- | --- | --- | --- | --- |
| **4AP** | $9.53 \pm 0.22$ | 7 | $-1.34 \pm 0.14$ | 5 | $3.08 \pm 0.07$ | 4 |
| **2,4DAP** | $9.09 \pm 0.03$ | 4 | $-2.01 \pm 0.10$ | 3 | $2.07 \pm 0.08$ | 4 |
| **3,4DAP** | $9.27 \pm 0.16$ | 4 | $-3.27 \pm 0.20$ | 3 | $1.91 \pm 0.09$ | 4 |
| **3Me4AP** | $9.75 \pm 0.26$ | 6 | $-1.27 \pm 0.08$ | 5 | $3.14 \pm 0.04$ | 4 |
| **4Me3AP** | $6.77 \pm 0.09$ | 6 | $0.07 \pm 0.06$ | 6 | $14.80 \pm 0.42$ | 4 |

\* Determined at pH 7.4.

*IC*_50_values for 3,4DAP (167 μ*M*) and 2,4DAP (821 μ*M*) were ∼1.7 greater and ∼2.8 lower, respectively. The *IC*_50_ determined for the methylated compound, 3Me4AP (43 μ*M*), was ∼7-fold greater than that produced by 4AP. Interestingly, the *IC*_50_of 4Me3AP (116 μ*M*) was ∼2.5-fold higher than that measured for 4AP. From the compounds measured, 4AP, 3,4DAP, and 3Me4AP had been measured before and our results are consistent with prior results^11,13^. On the other hand, compounds 2,4DAP, 4MeP, 2,4DMeP, 3,4DMeP, 4Me2AP and 4Me3AP have not been evaluated before. Our results indicate that 4MeP, 2,4DMeP, 3,4DMeP, and 4Me2AP are inactive (as expected given the lack of an exocyclic amino group). Our results also indicate that 4Me3AP is more potent than 4AP and 3,4DAP which indicates, for the first time, that having an amino group in the 4 position is not required for efficiently blocking K_V_ channels.

Next, we measured the values of the *pK*_*a*_, log *D* and *P*_*e*_of the active compounds. These measurements are presented in **Table I**. In terms of *pK*_*a*_ 4AP, 3,4DAP, 2,4DAP, and 3Me4AP had all values > 9 whereas 4Me3AP was close to 7. In terms of log *D* and *P*_*e*_ 4AP, 3,4DAP, 2,4DAP, and 3Me4AP were found to be moderately hydrophilic (log *D* < −1) and lowly permeable (*P*_*e*_ range 1.9 – 3.1 *cm*/*s*) whereas 4Me3AP was found to be significantly more lipophilic (log *D* ∼ 0.1) and permeable (*P*_*e*_=14.8 *cm*/*s*).

### pH and voltage dependence of the blocking of 4AP, 3Me4AP, and 4Me3AP

The canonical blocking mechanism of 4AP on Shaker K_V_ ion channel is a sequential process that is *pH*– and *V*-dependent^19,30^. Here, we evaluated the blockage of the Shaker K_V_ ion channel by the analogs 2,4DAP, 3,4DAP, 3Me4AP and 4Me3AP under different *pH* and *V* conditions. **Figure 3** summarizes these *pH*– and *V*-dependent behaviors. First, we determined the relative current curves assessed at different *pH* values (right panels) that ranged from 5.7 (or 6.8) to 9.1 and then the *IC*_50_*vs. V* curves (left panels) for 2,4DAP, 3,4DAP, 3Me4AP and 4Me3AP 4AP, respectively. With respect to the *pH* dependence, the trend observed of *IC*_50_determined for 2,4DAP, 4AP, 3,4DAP and 3Me4AP indicates that increasing the *pH* from 6.8 to 9.1 resulted in a decrease in the *IC*_50_, whereas *IC*_50_values of 4Me3AP exhibit an opposite trend and lower dependence on pH (**Figure 3d**). To analyze the variation of *IC*_50_with respect to *V* (*IC*_50_(*V*)) of these 4AP-based structural analogs, we fitted the data with the Hermann and Gorman equation (**Equation 2**) that describes binding mechanism of 4AP of K_V_ ion channel in terms of a simple of two state binding model^26^ and allows to estimate the electrical distance (δ) to be crossed by 4AP within the intense electric field generated in the channel to binds its site through the channel pore. These δ values are reported in **Table II** and ranged around 0.4 under all *pH* conditions evaluated in this study. These results suggest that the compounds share a common binding site and must traverse approximately half of the parametrized membrane electric field to reach it.

**Figure 3.**
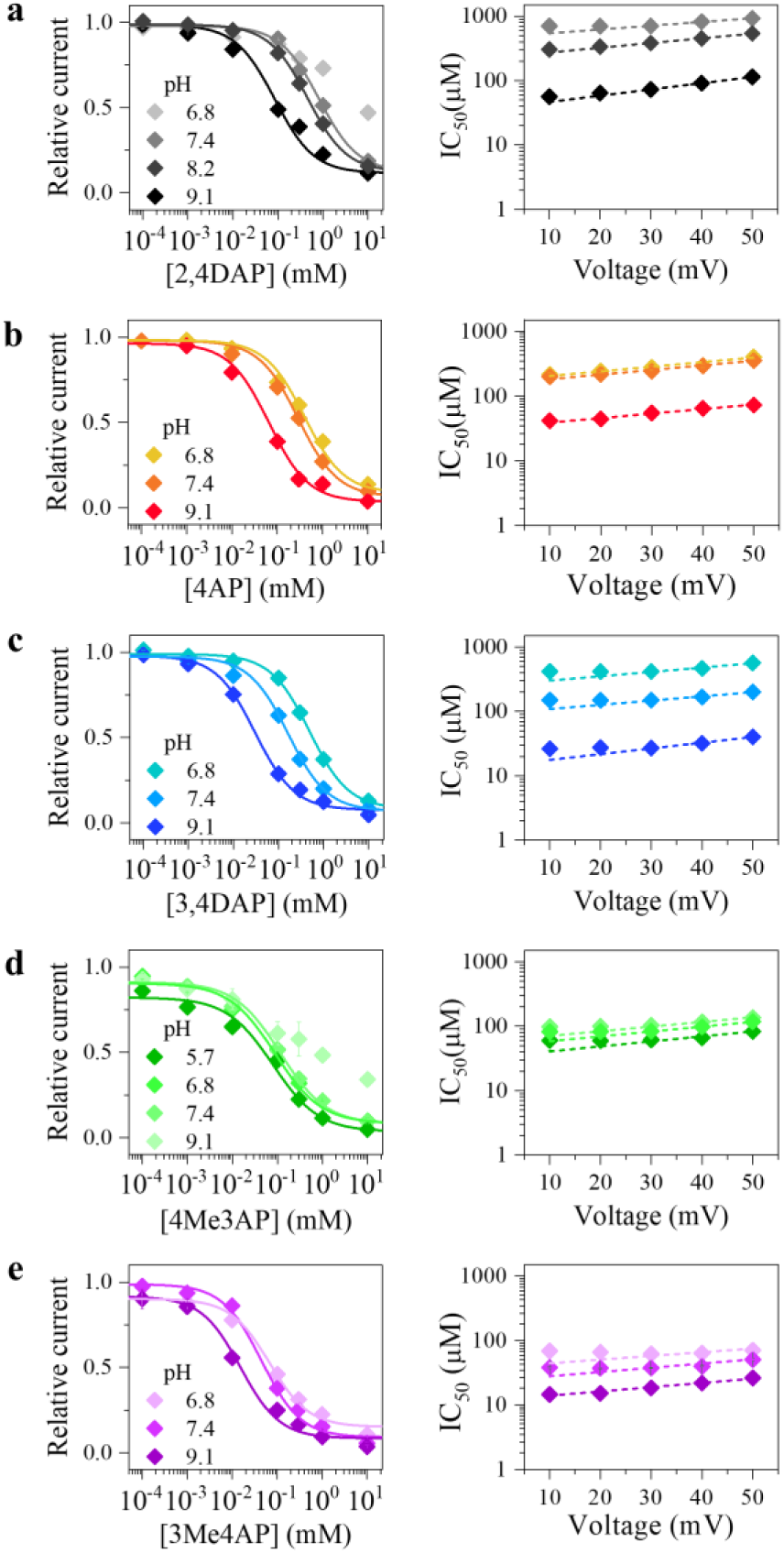
*pH*– and *V*-dependence of the blockade of the Shaker Kv channel by 4AP structural analogs. Relative current *vs.* concentration of 4AP-based structural analogs at several *pH* conditions and *V* = +40*mV* (left panels) and *IC*_50_*vs. V* curves (right panels) determined for: (**a**), 2,4DAP, (**b**), 4AP, (**c**), 3,4DAP, (**d**), 4Me3AP, and (**e**), 3Me4AP. Continuous lines in the left panels represent fits to the Hill equation (**Equation 1**) with a Hill coefficient ranging from 0.9 to 1.1. Hill fits parameters are shown in **Table I**. Dashed lines of the right panels represent the fits with the one-step *V*-dependent model of inhibition of Shaker K_V_ by these 4AP-based structural analogs (**Equation 2**), producing an electrical distance δ ∼ 0.4. Fits parameters of *IC*_50_*vs. V* curves are shown in **Table II**. Electrical recordings were obtained from *Xenopus laevis* oocytes expressing the Shaker K_V_ ion channel. Each structural analog was tested in independent oocytes, with the number of replicates ranging from 3 to 7. Electrical recordings are shown in the supplemental material (**Figure S2**).

**Table II.**
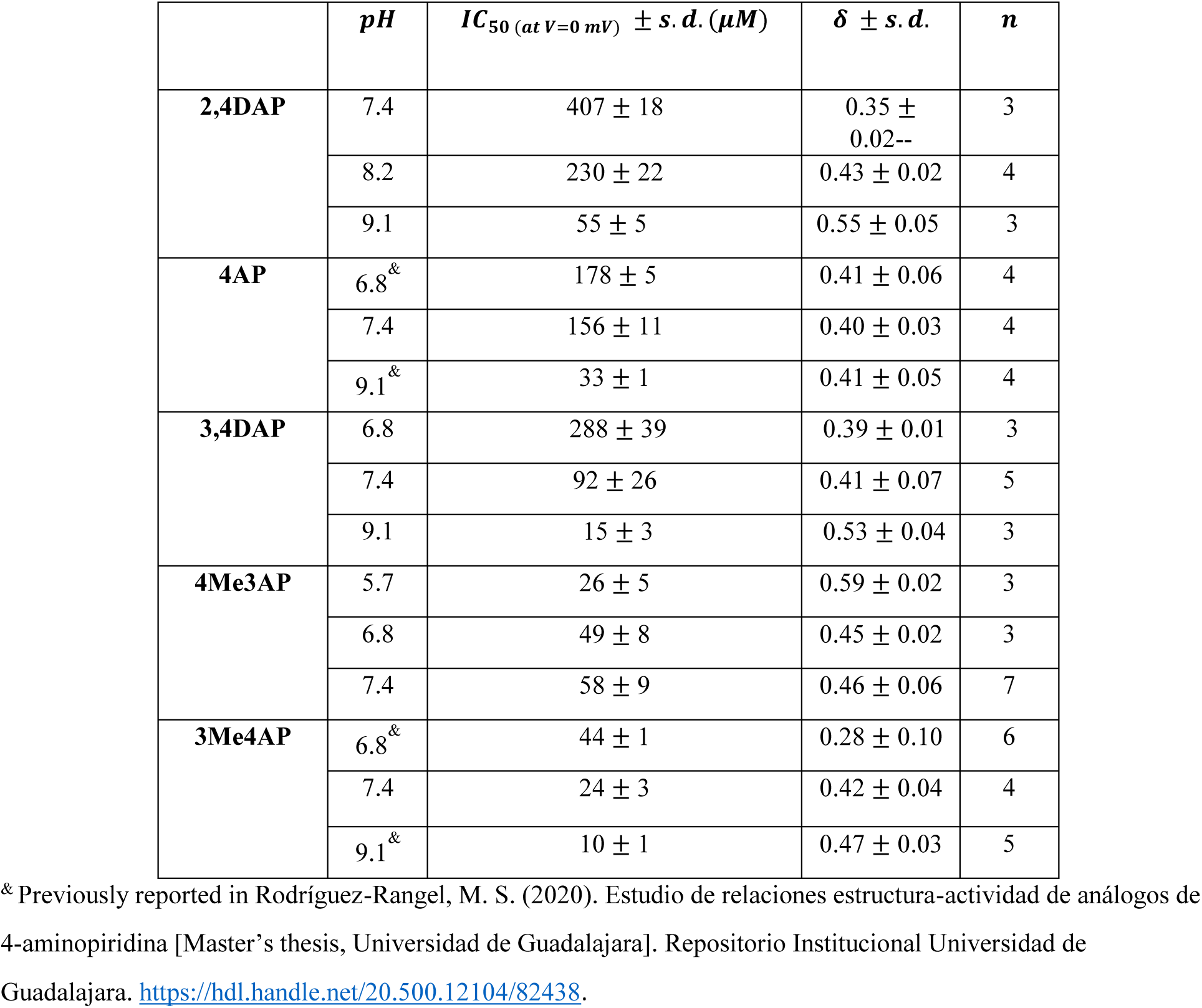
Fits parameters of the *IC*_50_(*V*) data with the Hermann-Gorman equation.

### Selective blockade of certain K_V_ ion channels expressed in the central nervous system and cardiac tissue by 4AP, 3Me4AP, and 4Me3AP

To begin to explore potential applications of these compounds, we evaluated the inhibition of specific human K_V_ ion channels expressed in the CNS (K_V_ 1.2, K_V_ 1.3, K_V_ 2.1, K_V_ 4.2, K_V_ 7.2, K_V_ 7.3, K_V_ 10.1 and the heart (Kv 11.1) by the molecules 4AP, 4Me3AP and 3Me4AP since these molecules exhibited the highest apparent affinity towards the canonical Shaker K_V_ ion channel from *drosophila* and also produced *pK*_*a*_, log *D* and *P*_*e*_ values (**Table I**) consistent with the ability to permeate lipid barriers such as the BBB. **Figure 4** shows representative K⁺ currents elicited from each K_V_ ion channel expressed in *Xenopus laevis* oocytes, recorded before and after the addition of 1 mM 4AP, 3Me4AP or 4Me3AP at *pH* 7.4 in response to a voltage step to +40 mV. As expected, 4Me3AP and 3Me4AP block the Shaker-related K_V_ 1.2 and K_V_ 1.3 ion channels, with 4Me3AP being the most potent (**Figure 4a** and **4b**). Interestingly, 4Me3AP molecule blocks the K_V_ 1.2 and K_V_ 1.3 ion channels with a higher apparent affinity than 3Me4AP and 4AP do (**Table III**). In contrast, 4AP, 4Me3AP, and 3Me4AP did not effectively block the Shab-related K_V_ 2.1 ion channel (**Figure 4c**). This result is consistent with previously reported results in the literature, as it is known that the *IC*_50_of 4AP for this channel is three orders of magnitude higher^18^. This pattern is also observed in the co-expressed K_V_ 2.1/6.4 channel, although compound 4Me3AP exhibits a higher apparent affinity than 4AP and 3Me4AP (**Figure 4d**).

**Figure 4.**
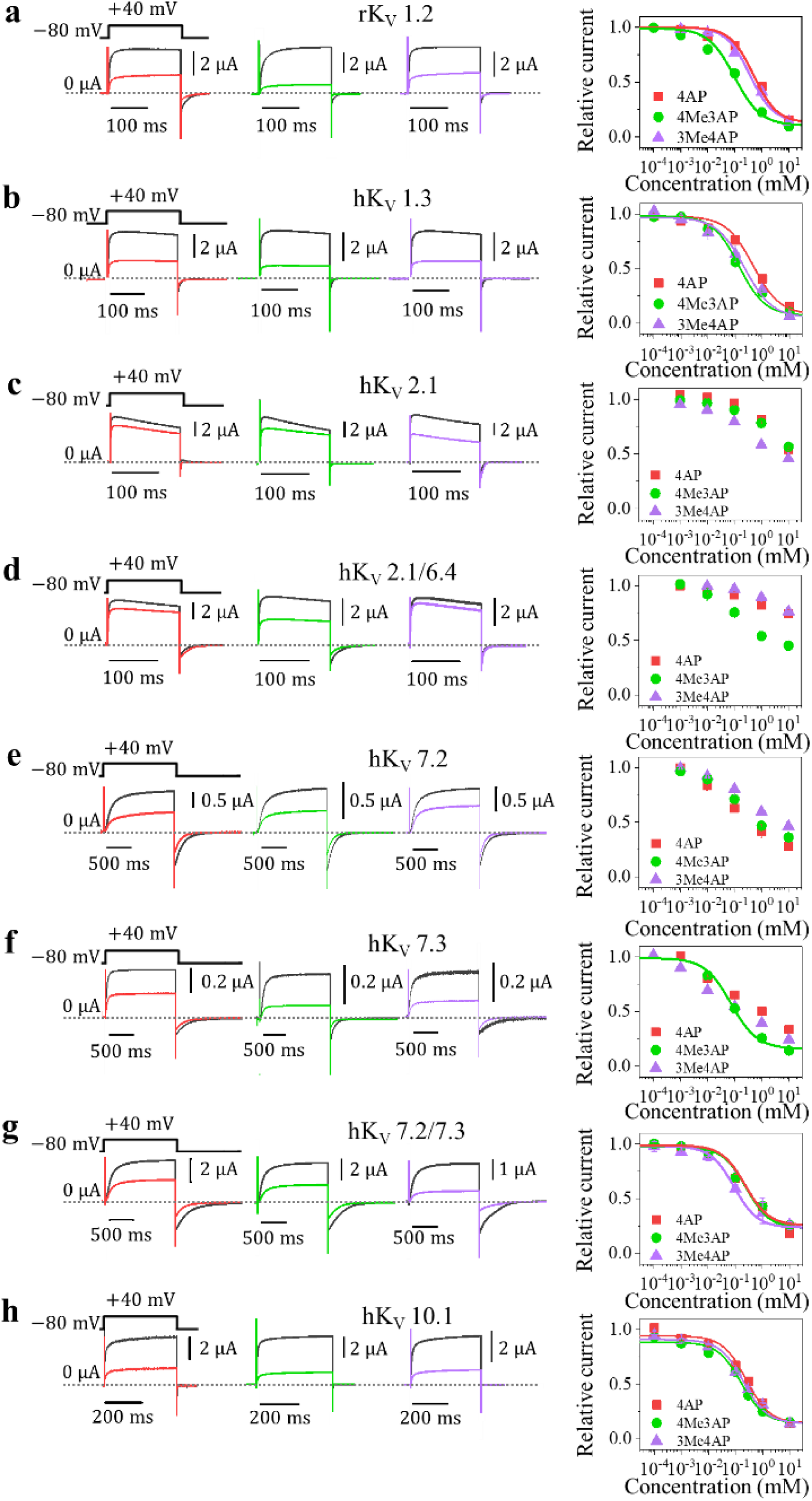
Specific blockage of certain K_V_ ion channels expressed in the SNC by 4AP, 3Me4AP and 4Me3AP. Representative recordings of K^+^ current elicited in response to a voltage stimulus (upper protocol) from different *Xenopus* oocytes expressing K_V_ ion channel commonly expressed in the SNC before (black) and after addition of 1 mM of 4AP (red), 4Me3AP (green), and 3Me4AP (purple) and relative current *vs.* concentration of 4AP, 4Me3AP, and 3Me4AP curves evaluated in (**a**), rK_V_ 1.2, (**b**), hK_V_ 1.3, (**c**), hK_V_ 2.1, (**d**), hK_V_ 2.1/6.4, (**e**), hK_V_ 7.2, (**f**), hK_V_ 7.3, (**g**), hK_V_ 7.2/7.3 and (**h**), hK_V_ 10.1 ion channels. Horizontal and vertical bars for each of K^+^ current recording indicate the time and current scale. Dashed lines represent the zero-current level. Continuous lines of the relative current curves represent the fits with the Hill equation (**Equation 1**) with a Hill coefficient ranging from 0.9 to 1.1. *IC*_50_ values from the Hill fits are shown in the **Table III**. The number of times each structural analog compound was tested in separate oocytes ranged from 3 to 7.

**Table III.** Estimated *IC*_50_ values at +40 *mV* and *pH* 7.4 of 4AP structural analogs assessed in different K_V_ ion Channels.

| Kv Ion Channel | Drug | *IC*<sub>50</sub>,<br>(95% C.I.)<br>(μM) | n |
| --- | --- | --- | --- |
| **1.2**<br>**(KCNA2)** | 4AP | 689, (546-832) | 3 |
|  | 3Me4AP | 398, (305-490) | 3 |
|  | 4Me3AP | 126, (100-150) | 3 |
| **1.3**<br>**(KCNA3)** | 4AP | 464 (402-526) | 3 |
|  | 3Me4AP | 319 (162-477) | 3 |
|  | 4Me3AP | 123 (66-181) | 3 |
| **7.3**<br>**(KCNQ3)** | 4AP | ** | 3 |
|  | 3Me4AP | ** | 1 |
|  | 4Me3AP | 141, (5-287) | 3 |
| **7.2/7.3**<br>**(KCNQ2/KCNQ3)** | 4AP | 289 (234-344) | 4 |
|  | 3Me4AP | 90 (74-99) | 2 |
|  | 4Me3AP | 288 (107-529) | 3 |
| **10.1**<br>**(EAG1)** | 4AP | 382 (334-432) | 4 |
|  | 3Me4AP | 181 (147-216) | 5 |
|  | 4Me3AP | 179 (147-210) | 7 |

We also examined the block of the K_V_ 7.2, K_V_ 7.3, and the co-expression K_V_ 7.2/K_V_ 7.3 ion channels by 4AP, 3Me4AP, and 4Me3AP (Figure 4e, 4f, and **4g**). Our results show that 4AP, 4Me3AP and 3Me4AP molecules inhibit ∼50% of the K^+^ currents generated by K_V_ 7.2 and K_V_ 7.3 at 1mM, however, the relative current curves could not be fitted using the Hill pharmacological model (**Equation 1**) with the exception of 4Me3AP for K_V_ 7.3 which yielded an *IC*_50_ of ∼141 μ*M* (**Figure 4f** and **Table III**). Unanticipatedly, relative current *vs.* concentration of 4AP, 3Me4AP and 4Me3AP from the co-expression of K_V_ 7.2/K_V_ 7.3 ion channels, produced a trend, from higher to lower apparent affinity, of 3Me4AP > 4Me3AP ≈ 4AP (**Figure 4g** and **Table III**). These findings confirm that the co-expression K_V_ 7.2/K_V_ 7.3 constitutes a fusion-protein-like motif expressed in neuronal nerves with different functional properties than K_V_ 7.2 and K_V_ 7.3^31^. Finally, our study was extended to include an analysis of the block of the K_V_ 10.1 ion channel by 4AP, 3Me4AP, and 4Me3AP (**Figure 4h**). Interestingly, our findings indicate that 3Me4AP and 4Me3AP block K_V_ 10.1 with ∼2-fold higher apparent efficiency than 4AP. These results demonstrate, for the first time, the inhibition of K_V_ 7.2, K_V_ 7.3, including the co-expression of K_V_ 7.2/K_V_ 7.3, and K_V_ 10.1 ion channels by 4AP, 3Me4AP, and 4Me3AP molecules.

It is well established that blockade of the K_V_ 11.1 ion channel is associated with severe ventricular arrhythmias, which may ultimately result in cardiac arrest^32^. For this reason, we evaluated the inhibitory effect of the novel K_V_ blocker molecule 4Me3AP on the Kv 11.1 channel expressed in Xenopus oocytes (**Figure 5**). **Figure 5a** and **5b** show representative recordings before and after the addition of 1 mM of 4AP, 3Me4AP, and 4Me3AP, and their relative current curves evaluated from 0.0001 to 10 mM, respectively. Our results show that, as is known for 4AP^33^, none of these compounds significantly blocks the heart channel.

**Figure 5.**
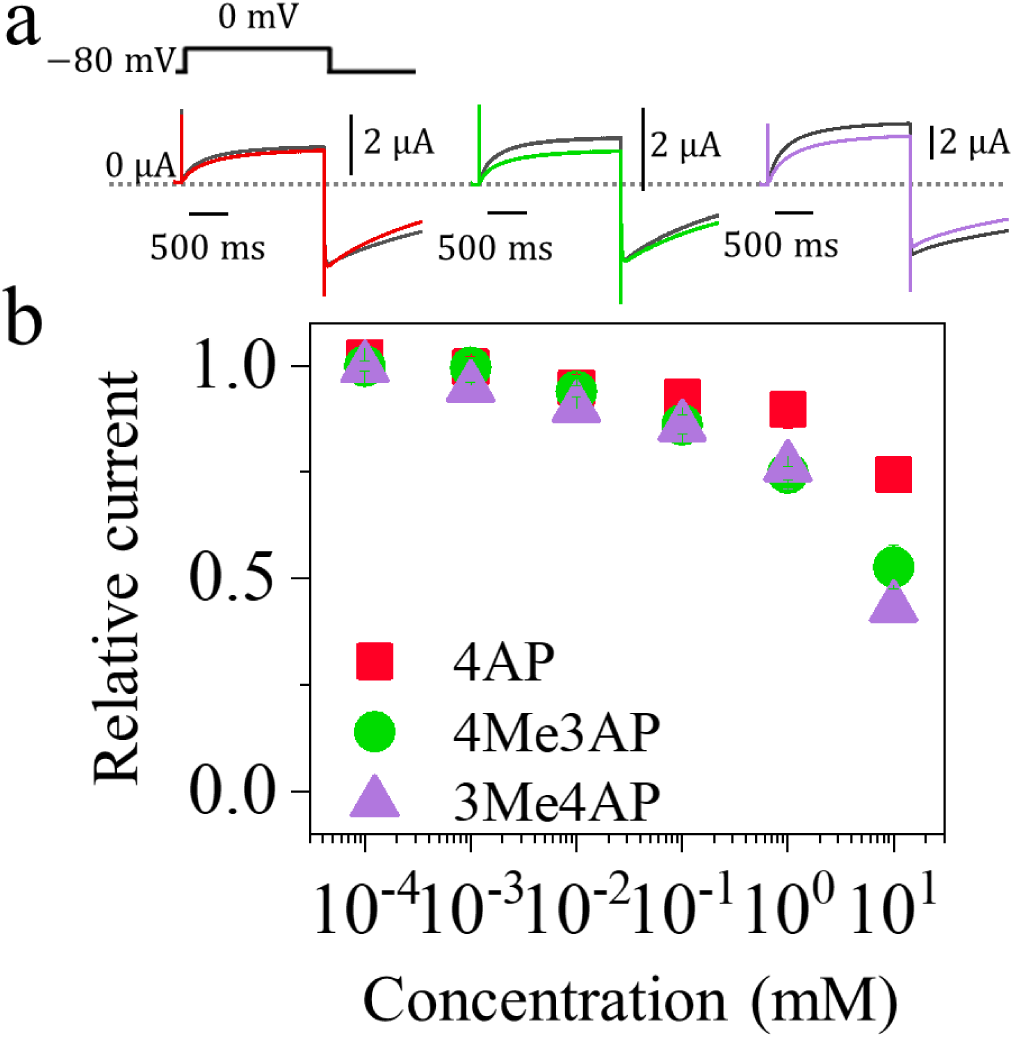
Block of Kv 11.1 ion channel by 4AP, 3Me4AP and 4Me3AP. (**a**), Representative K^+^ recordings elicited from three different oocytes expressing the K_V_ 11.1 ion channel as the response to a voltage stimulus protocol that consisted of 50 ms depolarization step from −80 to 0 mV (top left upper voltage protocol) before and after addition of 1 mM of 4AP, 4Me3AP and 3Me4AP. Dashed lines represent the zero-current level. (**b**), Relative current *vs.* the concentration of 4AP, 4Me3AP and 3Me4AP. Measurements were performed at *pH* 7.4.

### Acute toxicity of 4AP, 3Me4AP, and 4Me3AP

The acute toxicity of molecules 4AP, 3Me4AP, and 4Me3AP was evaluated in *BALB/c* mice following oral administration (**Table IV**). The three compounds produced comparable effects, such as tremors and seizures, which increased in both frequency and intensity with rising doses. The acute oral median lethal doses (*LD*_50_, in *mg*/*kg*) were 12.7, 13.23, and 29.3 for 4AP, 3Me4AP, and 4Me3AP, respectively. 4AP and 3Me4AP exhibited similar *LD*_50_values, whereas 4Me3AP showed a higher *LD*_50_, indicating lower toxicity in the *BALB/c* mice. This result suggests that 4Me3AP may possess a broader therapeutic margin compared with 4AP.

**Table IV.**
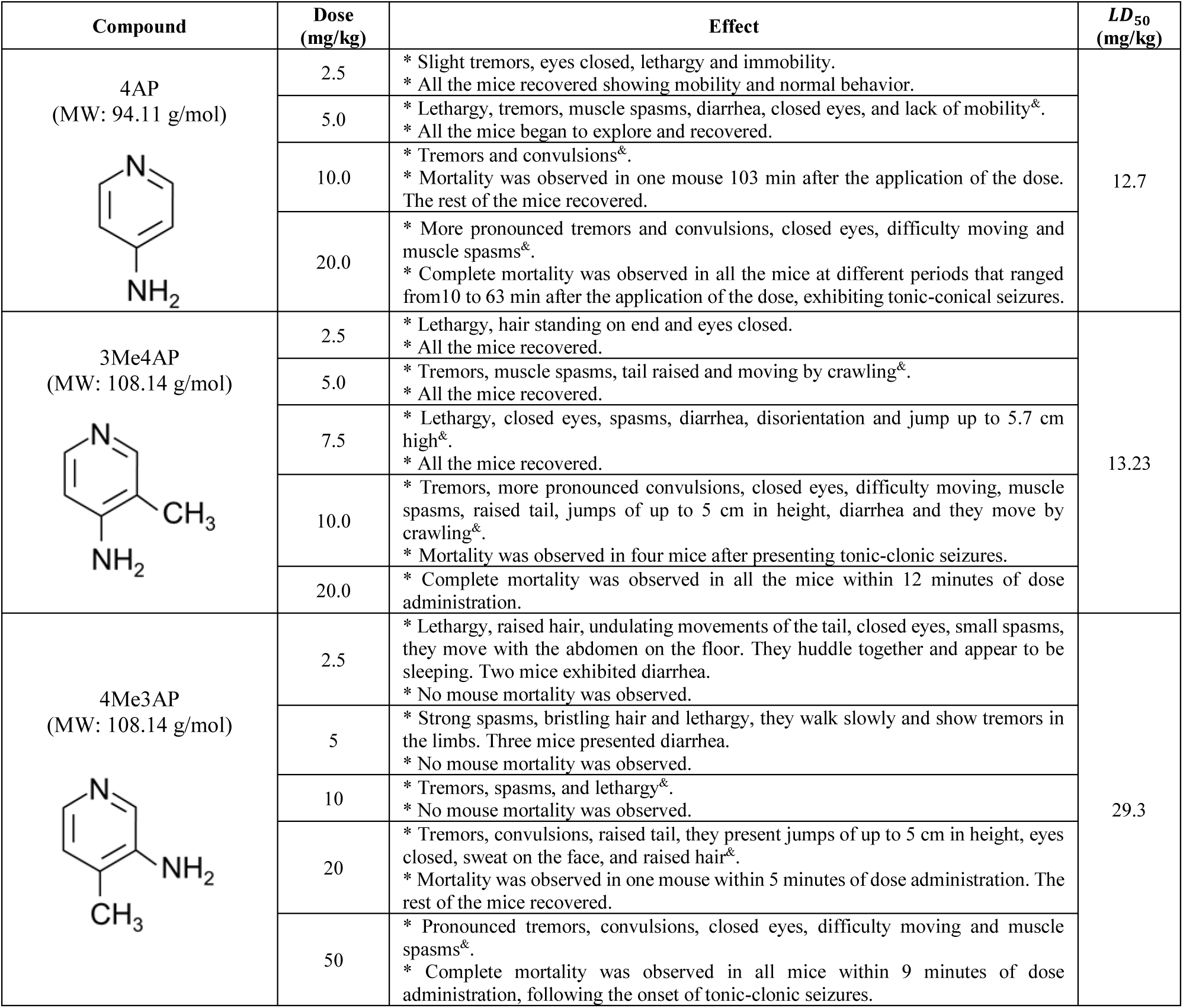
Acute toxicity study of 4AP, 4Me3AP and 3Me4AP in a murine model.

| Compound | Dose (mg/kg) | Effect | LD<sub>50</sub> (mg/kg) |
| --- | --- | --- | --- |
| **4AP**<br>(MW: 94.11 g/mol) | 2.5 | * Slight tremors, eyes closed, lethargy and immobility.<br>* All the mice recovered showing mobility and normal behavior. | 12.7 |
|  | 5.0 | * Lethargy, tremors, muscle spasms, diarrhea, closed eyes, and lack of mobility<sup>Ⓢ</sup>.<br>* All the mice began to explore and recovered. |  |
|  | 10.0 | * Tremors and convulsions<sup>Ⓢ</sup>.<br>* Mortality was observed in one mouse 103 min after the application of the dose. The rest of the mice recovered. |  |
|  | 20.0 | * More pronounced tremors and convulsions, closed eyes, difficulty moving and muscle spasms<sup>Ⓢ</sup>.<br>* Complete mortality was observed in all the mice at different periods that ranged from 10 to 63 min after the application of the dose, exhibiting tonic-clonic seizures. |  |
| **3Me4AP**<br>(MW: 108.14 g/mol) | 2.5 | * Lethargy, hair standing on end and eyes closed.<br>* All the mice recovered. | 13.23 |
|  | 5.0 | * Tremors, muscle spasms, tail raised and moving by crawling<sup>Ⓢ</sup>.<br>* All the mice recovered. |  |
|  | 7.5 | * Lethargy, closed eyes, spasms, diarrhea, disorientation and jump up to 5.7 cm high<sup>Ⓢ</sup>.<br>* All the mice recovered. |  |
|  | 10.0 | * Tremors, more pronounced convulsions, closed eyes, difficulty moving, muscle spasms, raised tail, jumps of up to 5 cm in height, diarrhea and they move by crawling<sup>Ⓢ</sup>.<br>* Mortality was observed in four mice after presenting tonic-clonic seizures. |  |
|  | 20.0 | * Complete mortality was observed in all the mice within 12 minutes of dose administration. |  |
| **4Me3AP**<br>(MW: 108.14 g/mol) | 2.5 | * Lethargy, raised hair, undulating movements of the tail, closed eyes, small spasms, they move with the abdomen on the floor. They huddle together and appear to be sleeping. Two mice exhibited diarrhea.<br>* No mouse mortality was observed. | 29.3 |
|  | 5 | * Strong spasms, bristling hair and lethargy, they walk slowly and show tremors in the limbs. Three mice presented diarrhea.<br>* No mouse mortality was observed. |  |
|  | 10 | * Tremors, spasms, and lethargy<sup>Ⓢ</sup>.<br>* No mouse mortality was observed. |  |
|  | 20 | * Tremors, convulsions, raised tail, they present jumps of up to 5 cm in height, eyes closed, sweat on the face, and raised hair<sup>Ⓢ</sup>.<br>* Mortality was observed in one mouse within 5 minutes of dose administration. The rest of the mice recovered. |  |
|  | 50 | * Pronounced tremors, convulsions, closed eyes, difficulty moving and muscle spasms<sup>Ⓢ</sup>.<br>* Complete mortality was observed in all mice within 9 minutes of dose administration, following the onset of tonic-clonic seizures. |  |

### Pharmacokinetics study of 4AP and 4Me3AP in a murine model

Pharmacokinetic profiles and parameters of 4AP and 4Me3AP in *BALB/c* mice are summarized in **Figure 6** and **Table V**. The pharmacokinetic curve of 4AP was consistent with a one-compartment model (**Figure 6a**), which assumes homogeneous distribution of the compound throughout the organism with a single phase of distribution and elimination. In contrast, the pharmacokinetic curve of 4Me3AP suggested a more complex pattern of absorption, distribution, and elimination. A two-compartment pharmacokinetic model (**Figure 6b**) appeared to describe the data trend best. Specifically, after oral administration, 4Me3AP may be rapidly distributed into a central compartment consisting primarily of blood circulation and highly perfused organs, followed by redistribution into a peripheral compartment likely composed of tissues, lipids, and organs with limited permeability barriers. In addition, the analysis of the pharmacokinetic curves indicated that under equivalent dosing conditions, 4Me3AP showed a two-fold higher *C*_*max*_compared with 4AP, indicating greater absorption and pharmacological potency. The *t*_*max*_of 4Me3AP and 4AP was around 10 *min*, reflecting similar absorption kinetics. The *t*_1/2_ of 4AP was comparable to the slow elimination phase of 4Me3AP (*t*_*e*1/2_ = 23.1 min). The *AUC*_0−250_of 4Me3AP was twice that of 4AP, consistent with higher bioavailability, while the clearance of 4Me3AP was half that of 4AP, indicating slower metabolism and/or elimination.

**Figure 6.**
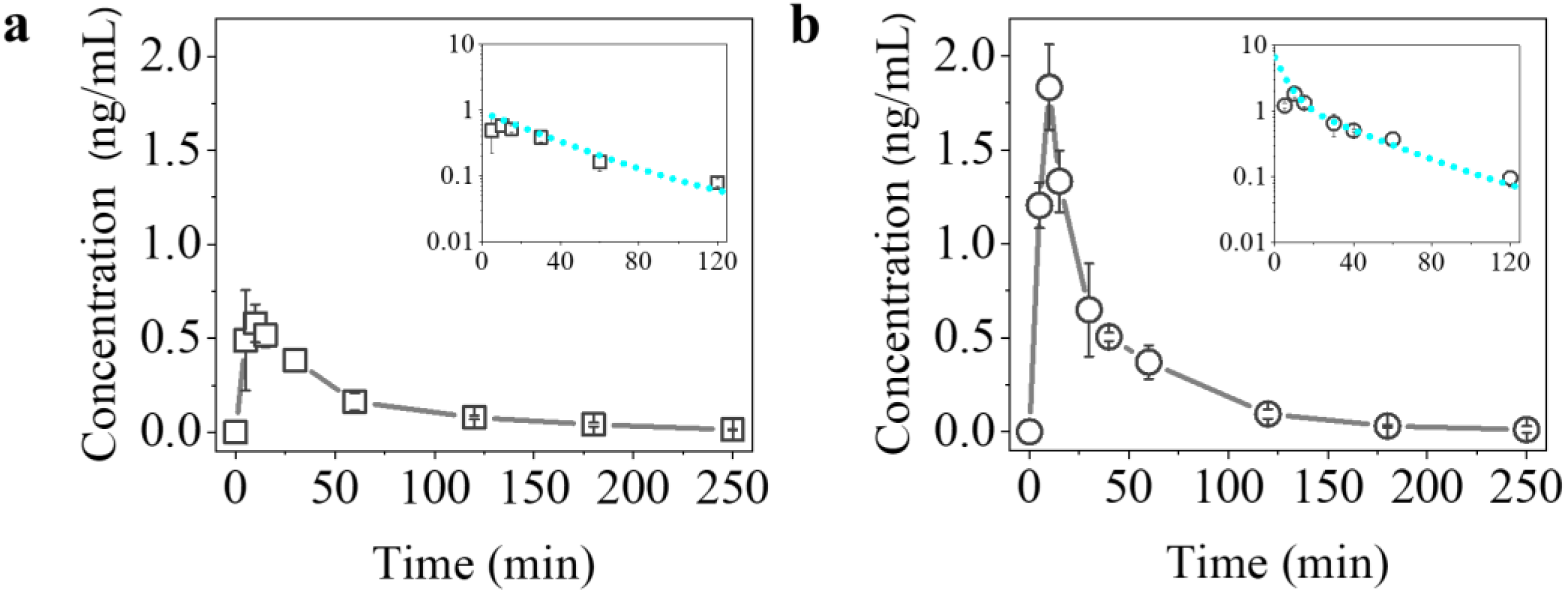
Single dose pharmacokinetic evaluation of 4AP and 4Me3AP in a murine model. **(a)**, Time course of plasma concentration of 4AP. (**b**), Time course of plasma concentration of 4Me3AP. Dashed cyan lines in the insets of panels a and b represent model fits to the elimination phase in plasma of 4AP or 4Me3AP using a one– or two-compartment pharmacokinetic model (**Equation 3**), respectively. The pharmacokinetic parameters obtained from the curve analysis are shown in **Table V**. Plasma concentrations of 4AP or 4Me3AP were measured at each time point in separate mice following a single dose (*see methods section*), and the number of independent measurements (*n*) ranged from 3 to 4.

**Table V.**
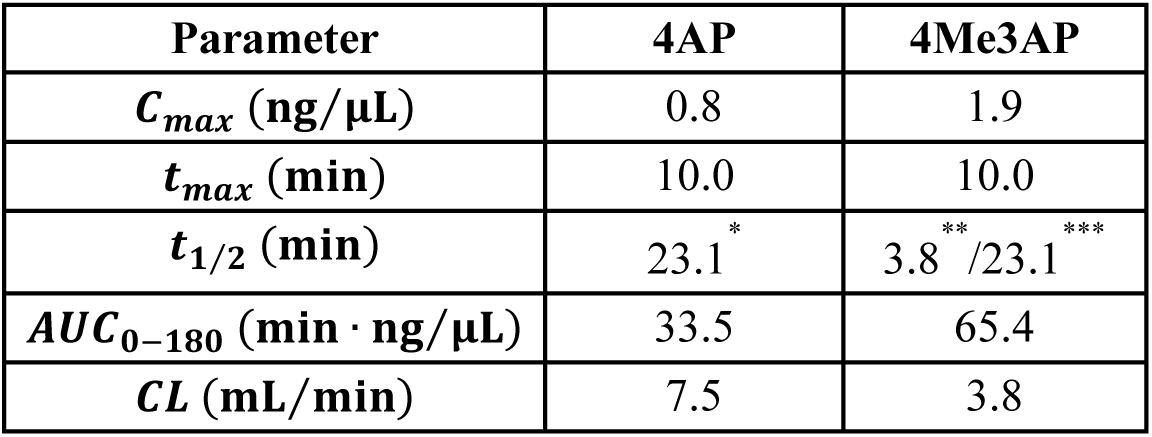
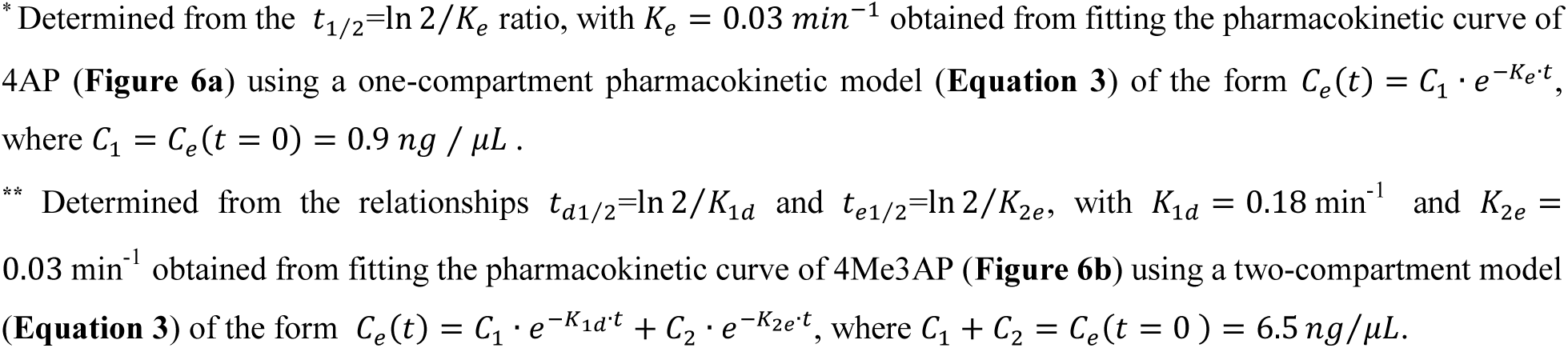
Pharmacokinetic parameters of 4AP and 4Me3AP.

| Parameter | 4AP | 4Me3AP |
| --- | --- | --- |
| $C_{max}$ (ng/μL) | 0.8 | 1.9 |
| $t_{max}$ (min) | 10.0 | 10.0 |
| $t_{1/2}$ (min) | 23.1<sup>*</sup> | 3.8<sup>**</sup> / 23.1<sup>***</sup> |
| $AUC_{0-180}$ (min · ng/μL) | 33.5 | 65.4 |
| $CL$ (mL/min) | 7.5 | 3.8 |

\* Determined from the $t_{1/2} = \ln 2 / K_e$ ratio, with $K_e = 0.03 \text{ min}^{-1}$ obtained from fitting the pharmacokinetic curve of 4AP (**Figure 6a**) using a one-compartment pharmacokinetic model (**Equation 3**) of the form $C_e(t) = C_1 \cdot e^{-K_e t}$ , where $C_1 = C_e(t = 0) = 0.9 \text{ ng} / \mu\text{L}$ .
\*\* Determined from the relationships $t_{d1/2} = \ln 2 / K_{1d}$ and $t_{e1/2} = \ln 2 / K_{2e}$ , with $K_{1d} = 0.18 \text{ min}^{-1}$ and $K_{2e} = 0.03 \text{ min}^{-1}$ obtained from fitting the pharmacokinetic curve of 4Me3AP (**Figure 6b**) using a two-compartment model (**Equation 3**) of the form $C_e(t) = C_1 \cdot e^{-K_{1d} t} + C_2 \cdot e^{-K_{2e} t}$ , where $C_1 + C_2 = C_e(t = 0) = 6.5 \text{ ng} / \mu\text{L}$ .

## Discussion

Aminopyridines have long been recognized for their therapeutic efficacy in neurological disorders involving impaired synaptic transmission or disrupted axonal conduction. Clinically, 4-aminopyridine (4AP) and 3,4-diaminopyridine (3,4DAP) improve neurological function in conditions such as multiple sclerosis and Lambert–Eaton myasthenic syndrome by enhancing conduction and neurotransmitter release^1–9^. More recently, radiolabeled aminopyridines, including [^18^F]3F4AP, [^11^C]3Me4AP, [^11^C]3CF_3_4AP, [^11^C]3MeO4AP, and [^18^F]5Me3F4AP, have emerged as novel imaging agents capable of detecting demyelinated axons *in vivo* by binding K_v_ channels that become exposed and upregulated after myelin loss^11,22,23,34–37^. These therapeutic and imaging applications share the same underlying principles governing aminopyridine pharmacology: membrane permeability, protonation state, voltage-dependent binding, and interactions with the intracellular cavity of K_v_ channels. Our findings extend this framework by demonstrating that 3-amino-4-methylpyridine (4Me3AP), the first characterized compound derived from a 3-aminopyridine scaffold, exhibits robust K_v_-blocking activity and favorable pharmacological properties, challenging the long-standing assumption that effective aminopyridine blockers must be based on the 4AP scaffold with an amino group fixed at the 4-position.

Mechanistically, 4Me3AP behaves in a manner consistent with classical aminopyridines. These molecules exist in a *pH*-dependent equilibrium between protonated and neutral species, determined by their *pK*_*a*_; the neutral form permeates lipid membranes, including the blood–brain barrier, while the protonated form is required for binding and blocking K_V_ channels from the intracellular side. Our *pH*– and *V*-dependence studies show that 4Me3AP must traverse approximately half of the membrane electric field to reach its binding site, much like 4-AP and related analogues, supporting the idea of a shared intracellular binding locus within KCNA channels. Although these data strongly suggest a common mechanism, structural, computational, and mutagenesis studies will be needed to definitively map the binding pose of 4Me3AP.

The structure–activity relationships (SAR) observed for 4Me3AP further parallel established aminopyridine trends. Incorporation of the amino group in the 3-position decreases basicity compared to 4AP, and methyl substitution on the 4-position increases lipophilicity. These changes increase membrane permeability and potency, which may be desirable pharmacological features for CNS therapeutic applications.

Overall, this study demonstrates that 4Me3AP possesses strong Kv-blocking activity and favorable physicochemical and pharmacokinetic properties, making it a promising molecule for further therapeutic exploration. While the present work examines only a single derivative and cannot support broad conclusions about the 3-aminopyridine scaffold as a whole, it provides the first evidence that 3-aminopyridine–based compounds can retain the key mechanistic features required for effective K_V_ inhibition. By challenging assumptions about positional requirements on the aminopyridine scaffold, these findings open new avenues for the rational design of next-generation aminopyridines tailored for therapeutic use and, potentially, future imaging applications.

## Acknowledgements

We thank Leon Islas (UNAM) for the rK_V_1.2 plasmid; Francisco Bezanilla (University of Chicago) for the hK_V_1.3 plasmid; Pablo Miranda (NINDS, NIH) for the hK_V_2.1 and hK_V_ 6.5 plasmids; and Carlos Villalva (University of the Pacific) for the hK_V_7.2 and hK_V_7.3 plasmids. hK_V_ 11.1 (pSP64-hERG1a) was a gift from Michael Sanguinetti (University of Utah) and K_V_ 10.1 was a gift from Luis Pardo (Max Planck Institute for Multidisciplinary Sciences). J.E.S.R. received financial support from the Technology Transfer Office of the Universidad de Guadalajara for this study. S.R.R. was supported by a fellowship from CONACyT (currently known as SECIHTI), Mexico (886951). PB received support from NIH/NINDS grant R01NS114066.

## Author contributions

S.R.R.: Designed research, performed the COVC, *pK*_*a*_and log *D* experiments, analyzed the data and contributed with the optimization of the HPLC method for the pharmacokinetics assays and performed pharmacokinetics experiments. O.G.C. and B.M.O: Optimized the HPLC method, performed the acute toxicology and pharmacokinetics experiments and analyzed the data. Y.S., S.E.S. and P.B. conducted *P*_*e*_experiments. P.B. also contributed to research design, data interpretation, co-wrote the manuscript and secured funding. J.E.S.R.: designed research, performed *pK*_*a*_and log *D* experiments and analyzed data, conceived and supervised the project, data interpretation, wrote the manuscript and secured funding. All authors revised and approved the final version of the manuscript.

## Conflict of Interest

An international PCT patent application related to the therapeutic use of 4Me3AP and related compounds has been filed, listing J.E.S.R. and S.R.R. as inventors. Additional patent applications have been filed related to the use of other aminopyridine derivatives for PET imaging and therapy with J.E.S.R., S.R.R., Y.S., and P.B. as inventors. The other authors declared no conflict of interest.

## Data Availability

The datasets generated during the current study are available from the corresponding author upon reasonable request.

